# Molecular determinants underlying DS2 activity at δ-containing GABA_A_ receptors

**DOI:** 10.1101/2021.01.21.427670

**Authors:** Christina B. Falk-Petersen, Frederik Rostrup, Rebekka Löffler, Stine Buchleithner, Kasper Harpsøe, David E. Gloriam, Bente Frølund, Petrine Wellendorph

## Abstract

Delta selective compound 2 (DS2) is one of the most widely used tools to study selective actions mediated by δ subunit-containing GABA_A_ receptors. DS2 was discovered over 10 years ago, but despite great efforts, the precise molecular site of action has remained elusive.

Using a combination of computational modeling, site-directed mutagenesis and cell-based pharmacological assays, we probed three potential binding sites for DS2 and analogs at α_4_β_1_δ receptors: an α_4_^(+)^δ^(-)^ interface site in the extracellular domain (ECD), equivalent to the diazepam binding site in αβγ_2_ receptors, and two sites in the transmembrane domain (TMD); one in the α_4_^(+)^β_1_^(-)^ and one in the α_4_^(-)^β_1_^(+)^ interface, with the α_4_^(-)^β_1_^(+)^ site corresponding to the binding site for etomidate and a recently disclosed low-affinity binding site for diazepam. We show that mutations in the ECD site did not abrogate DS2 modulation. However, mutations in the TMD α_4_^(+)^β_1_^(-)^ interface, either α_4_(S303L) of the α_4_^(+)^-side or β_1_(I289Q) of the β_1_^(-)^-side, convincingly disrupted the positive allosteric modulation by DS2. This was consistently demonstrated both in an assay measuring membrane potential changes and by whole-cell patchclamp electrophysiology and rationalized by docking studies. Importantly, general sensitivity to modulators was not compromised in the mutated receptors. This study sheds important light on the long-sought molecular recognition site for DS2, refutes the misconception that the selectivity of DS2 for δ-containing receptors is caused by a direct interaction with the δ-subunit, and instead points towards a functional selectivity of DS2 and its analogs via a surprisingly well-conserved binding pocket in the TMD.

**Significance statement:** δ-Containing GABA_A_ receptors represent potential drug targets for the treatment of several neurological conditions with aberrant tonic inhibition. Yet, no drugs are currently in clinical use. With the identification of the molecular determinants responsible for positive modulation by the know compound DS2, the ground is laid for design of ligands that selectively target δ-containing GABA_A_ receptor subtypes, for better understanding of tonic inhibition, and, ultimately, for rational development of novel drugs.

## Introduction

Inhibition in the brain is primarily mediated by g-aminobutyric acid (GABA) acting through GABA receptors, with the ionotropic GABA_A_ receptors (GABA_A_R) being responsible for fast inhibition. Thus, GABA_A_Rs play an essential role in transmitting inhibitory signaling in the brain. Structurally speaking, GABA_A_Rs belong to the Cys-loop receptor family of pentameric receptor complexes and are composed from a repertoire of 19 different subunits in mammals, with the most commonly expressed in the CNS being α_1-6_, β_1-3_,γ_1-3_ and δ (1). The subunit stoichiometry of the archetypical GABA_A_ receptor is 2α, 2β and a third subunit most typically being either a γ or a δ-subunit, but other stoichiometries have also been reported (1). Studies on the subunit arrangement of the most abundantly expressed synaptic subtype, α_1_β_2_γ_2_, and a number of other γ-containing subtypes, show that the subunits are arranged as γ-β-α-β-α in a counterclockwise fashion around the central ion channel (2, 3). Although it is generally accepted that the δ-subunit in its cognate receptors simply replaces the g-subunit with respect to arrangement (4), this is still not unequivocally established (5–8). Irrespectively, the orthosteric binding sites are located at the β^(+)^α^(-)^ interfaces in the extracellular domain (ECD), and a number of allosteric binding sites have also been identified in the subunit interfaces in both the ECD and transmembrane domain (TMD) (9). These include for example the benzodiazepine site in the ECD α^(+)^γ^(-)^ interface, responsible for mediating the anxiolytic and sleep-inducing effect of the benzodiazepines, including diazepam (Valium^®^), widely used in the clinic (10, 11).

The δ-containing GABAaRs are located primarily at extrasynaptic sites where they mediate tonic (persistent) inhibition (12, 13), hence controlling neuronal excitability (14). Tonic inhibition is involved in various physiological responses and pathophysiological conditions (15) underlining δ-GABA_A_R as valuable drug targets. THIP (gaboxadol), a δ-selective super agonist (16), was in a clinical phase III trial for treatment of primary insomnia, but failed (17) and have been in clinical trials (OV101) for the treatment of two rare disorders Fragile-X (trial number NCT03697161) (18) and Angelman syndrome (trial number NCT02996305) (19). Additionally, several neurological disorders such as stroke (20, 21) and epilepsy (22, 23), underline the continued interest in targeting these receptors. This also includes an emerging relevance for immunomodulation via actions on peripheral δ-containing receptors or equivalents (24, 25). However, compared to the synaptic γ-containing receptors, pronounced insight into the physiological and pathophysiological role of δ-containing receptors is still limited by the low number of potent and selective compounds.

In addition to THIP, another compound with selectivity for the δ-containing receptors is the positive allosteric modulator (PAM) delta selective compound 2 (DS2; 4-Chloro-*N*-[2-(2-thienyl)imidazo[1,2-*a*]pyridin-3-yl]benzamide) (26). DS2 has been used extensively as a tool compound to confirm the presence of δ-receptor-mediated tonic currents both in *in vitro* and *in vivo* studies (27–30). DS2 was identified in a screening campaign and reported as a δ-selective PAM at α_4_β_3_δ GABA_A_Rs, showing no or limited effects at α_4_β_3_γ_2_ and α_1_β_3_γ_2_ receptors (26). This selectivity was confirmed in thalamic relay neurons, where only extrasynaptic tonic currents were enhanced (26). In 2013, Jensen *et al*. confirmed the reported selectivity for δ-containing receptors, using δ^-/-^ mice showing that this effect is clearly dependent on the δ-subunit (31). DS2 displays limited brain permeability (31) but was, nonetheless, shown to improve recovery after stroke in mice, plausibly by dampening peripheral immune activation (24). Recently, a methoxy analog of DS2, termed DS2OMe, was identified and confirmed to have a similar potency and subtype selectivity as DS2. Interestingly, the derived positron emission tomography (PET) tracer of DS2OMe, [^11^C]DS2OMe, has been shown to have potential as a PET tracer for visualization of δ-containing receptors in brains of larger mammals such as pig (32).

In 2018, the first cryogenic electron microscopy (cryo-EM) structure of a human GABA_A_R pentamer α_1_β_2_γ_2_ was published (33). Following this, the structure of the human α_1_β_3_γ_2_ receptor was solved in complex with diazepam, revealing both the known high-affinity diazepam binding site in the α^(+)^γ^(-)^ interface in the extracellular domain (ECD), and a novel low affinity binding site located in the α^(+)^β^(-)^ interface of the transmembrane domain (TMD) of the receptor (34). This low-affinity diazepam binding site is located in the same pocket as the binding site for the general anesthetics (e.g. etomidate) (35) which is also the proposed binding site for AA29504 (36). Additionally, similar sites are suggested to be present in all five inter-subunit interfaces in the TMD (37–39).

Based on the notion that binding pockets evolved through nature are often highly conserved, combined with the structural similarities between DS2 and benzodiazepine ligand zolpidem (40), we hypothesized that these diazepam binding pockets are similarly present in δ-containing subtypes and that either of them could represent the long-sought DS2 molecular recognition site for DS2.

We here report the identification of two residues, α_4_(S303) and β_1_(I289), within the predicted α_4_^(+)^β_1_^(-)^ interface of the TMD in α_4_β_1_δ receptors, that are necessary for DS2 modulation. This is supported by docking of DS2 and analogs into the identified binding pocket.

## Material and Methods

### Chemicals and materials

The compounds DS2; (4-chloro-N-[2-(2-thienyl)imidazo[1,2-a]pyridin-3-yl]benzamide), AA29504; ([2-amino-4-(2,4,6-trimethylbenzylamino)-phenyl]-carbamic acid ethyl ester), etomidate; ((*R*)-1-(1-phenylethyl)-1H-imidazole-5-carboxylic acid ethyl ester), picrotoxin and GABA were obtained from Tocris Bioscience (Bristol, UK). DS2OMe (4-Methoxy-*N*-[2-(thiopen-2-yl)imidazole[1,2-*a*]pyridine-3-yl]benbamide) was synthesized in-house as described previously (32). DMEM with GlutaMAX-I, FBS, penicillin-streptomycin, hygromycin B, trypsin-EDTA, DPBS and HBSS were purchased from Life Technologies (Paisley, UK). DMSO, HEPES, MgCl_2_, CaCl_2_, poly-D-lysin (PDL) and MgATP were purchased from Sigma-Aldrich (St. Louis, MO, USA). The FLIPR membrane potential Blue dye was purchased from Molecular Devices (Crawley, UK) and Polyfect transfection reagent from Qiagen (West Sussex, UK). Stocks of DS2 and DS2OMe were prepared at 1 mM and 10 mM concentrations in DMSO with final DMSO concentration <0.1%. Due to moderate solubility and 4x concentrations used in the FMP assay, the FMP buffer was preheated to 37 °C in a water bath before addition of compound and preparation of serial dilutions. Only stocks with final concentrations below 12 μM were used for further dilutions. Furthermore, higher concentrations were prepared separately.

### Cells and transfections

A HEK-293 Flp-In cell line stably expressing the human δ-GABA_A_R subunit (28) was used for transfection with human α- and β-subunits to express recombinant wildtype and mutant GABA_A_Rs and transfection ratios optimized as described (28). Cells were maintained in DMEM containing GlutaMAX-I, supplemented with 10% FBS and 1% penicillin-streptomycin and kept in an incubator at 37 °C and a humidity of 5% CO_2_. 200 μL/ml hygromycin B was added to the media as positive selection. Transfection was performed using Polyfect (Qiagen) following the manufacturer’s instructions except for using half the volume of transfection reagent for each transfection. α- and β-subunits were co-transfected in a 1:1 ratio for FMP experiments, and for patch-clamp experiments additionally co-transfected with GFP in a 0.5:1:1 ratio (0.8 μg:1.6 μg:1.6 μg in 6 cm culture dishes) in order to visualize transfected cells.

### Plasmids and mutant constructs

The plasmids used for transfection to transiently express GABA_A_ receptors have been described previously (28). The WT human α_4_ and β_1_ subunits were subcloned into the pUNIV vector (Addgene, Cambridge, MA, USA) and the human δ-subunit into the pcDNA5/FRT vector (Invitrogen, Paisley, UK) using the δ-construct described previously (28). Plasmids carrying single and double mutations were generated and sequence-verified by GenScript (Piscataway, NJ, USA). The numbering of the mutants refers to the sequences with the signal peptide included.

### Generation of stable cell lines

Mutations introduced into the δ-subunit were established as stable HEK293 Flp-In^TM^ cell lines (Invitrogen, Paisley, UK), generating a stable cell line for each mutant. The stable cell lines were generated using the pcDNA/FRT/V5-His TOPO TA Expression kit (Invitrogen) performed according to the manufacturer’s protocol and as described previously (28), except for using 25 μL Polyfect and 4 μg DNA for transfection in a 10 cm culture dish.

### FLIPR membrane potential (FMP) assay

The FLIPR membrane potential (FMP) assay was performed exactly as described previously (28). In brief, 48 hours before the assay, cells were transfected. 16-20 hours later, cells were plated into clear-bottomed PDL (poly-D-lysine)-coated black 96-well plates in a number of 50,000 cells/well, suspended in cell media, and placed in an incubator at 37 °C with a humidity of 5% CO_2_ until performing the assay. 44-48 hours post-transfection, the media was removed, cells were washed in assay buffer (100 μL/well) and incubated in 100 μL/well 0.5 mg/mL FMP blue dye freshly dissolved in assay buffer (HBSS containing 20 mM HEPES adjusted to pH 7.4 and supplemented with 2 mM CaCl_2_ and 0.5 mM MgCl_2_) for 30 min shielded from light in an incubator at 37° C and a humidity of 5% CO2. Ligand solutions were prepared in 4x assay buffer and added to a ligand plate, which was placed in the FLEXstation3 plate reader (Molecular Devices, Crawley, UK), preheated to 37° C for temperature equilibration for 10-15 min. After transferring the cell plate to the reader, the fluorescence was measured at baseline and after ligand addition by detecting emission at 560 nm caused by excitation at 530 nm.

### FMP data analysis, exclusion criteria and statistics

Compound-induced signals were reported as changes in fluorescence units (ΔRFU), with the signal given as the difference between the average of the baseline signal (approx. 30 s recording) subtracted the peak response (or minimum response for decreases in baseline). All raw traces were manually inspected for obvious artefacts after compound addition. For high concentrations of DS2 (1-20 μM) we regularly observed negative RFU values below the buffer responses that in certain cases were excluded (see below). This phenomenon was independent of receptor subtype as it was observed for both δ-HEK and mock cells. The phenomenon was less pronounced for DS2OMe why this compound was preferred in some sub-studies. To circumvent this problem, we set up the following exclusion criteria: negative ΔRFU or decreased ΔRFU values for high concentration (>1 μM) of DS2 and DS2OMe compared to the ΔRFU for a lower concentration in the same experiment (indicative of precipitation). Additionally, as a rule of thumb, the group size was to be at least n=3, or what was needed to obtain a SEM value < 0.2. Additionally, curve fittings resulting in ambiguous EC_50_ values, and R^2^ values lower than 0.80 were omitted from analyses.

Concentration-response curves were fitted using nonlinear regression, with the log-transformed concentration as x-values, using the four-parameter concentration-response equation:

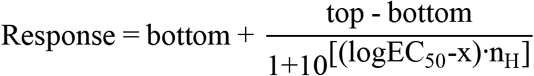

to determine the EC_50_ value and Hill slope. The ‘bottom’ and ‘top’ denotes the upper and lower plateau of the curve, respectively. The calculated EC_50_ values were log-transformed to obtain mean pEC_50_ values. Statistical analysis of mutated receptors was performed on the pEC_50_ values using the two-sided Welch’s t-test compared to WT, correcting for multiple comparison using the original FDR method of Benjamini and Hochberg with a discovery rate of 0.05. Both adjusted and un-adjusted P-values are reported. The data and statistical analysis were performed in GraphPad Prism (v. 8.4.3; GraphPad, San Diego, CA, USA).

### Whole-cell patch-clamp electrophysiology

Whole-cell patch-clamp experiments were performed on δ-HEK cells transiently co-expressing WT or mutant α- and β-subunits and GFP as described previously (41). In short, the transfected cells were transferred to 35 mm petri dishes (100,000-200,000 cells) the day prior to performing the experiment. On the day of experiment, cell media was exchanged for ABSS (containing the following (in mM): NaCl 140, KCl 3.5, Na_2_HPO_4_ 1.25, MgSO_4_ 2, CaCl_2_ 2, glucose 10, and HEPES 10; pH 7.35) at room temperature (20-24 °C), before placing at the stage of an Axiovert 10 microscope (Zeiss, Germany). Viewing the cells at 200× magnification and visualizing cells containing green fluorescent protein with UV light from an HBO 50 lamp (Zeiss, Germany), the cells were approached with micropipettes of 1.2-3.3 MΩ resistance manufactured from 1.5 mm OD glass (World Precision Instruments, Sarasota, Florida, USA) on a microelectrode puller, model PP-830 (Narishige, Tokyo, Japan). The micropipettes contained an intracellular solution composed of the following (in mM): KCl 140, MgCl_2_ 1, CaCl_2_ 1, EGTA 10, MgATP 2, and HEPES 10; pH 7.3.

Recordings were made from cells in the whole-cell configuration using the standard patchclamp technique in voltage mode and an EPC-9 amplifier (HEKA, Lambrecht, Germany). The clamping potential was −60 mv and series resistance was 80% compensated. Whole-cell currents were recorded using Pulse and PulseFit software (v.8.80, HEKA). Ligand solutions, prepared in ABSS, were applied using two VC3-8xP pressurized application systems feeding into a sixteen-barreled perfusion pipette (ALA Scientific Instruments Inc., Farmingdale, NY, USA) ending approximately 100 μm from the recorded cell. PAMs were tested using coapplication with a concentration of GABA corresponding to GABA EC_10-35_ at the respective receptor subtype. Preapplication was not used for DS2 and DS2OMe, as results from preliminary experiments showed no difference in the size of the peak current with and without preapplication of the PAM. PAMs and GABA was co-applied for 10-30 s until the peak current was reached. Agonists was applied for 5 s. Between compound applications, compound-free ABSS was applied from one of the barrels in order to quickly remove the compounds from the cell and cells were allowed to recover for 1 min before the next ligand application.

### Patch-clamp data analysis and statistics

All currents were normalized to the maximum GABA current and given as % I/I_max_. All currents are reported as normalized mean currents with 95% confidence interval, determined based on currents from at least five different cells from at least two transfections. Datasets with GABA controls (0.1-0.5 μM) deviating from GABA EC_10_ to EC_35_ were excluded from the analysis.

Statistical analysis was applied to test whether the PAMs potentiated the GABA control response using two-side Welch’s t-test as for FMP data. Analysis of currents was performed using Pulse and Pulsefit (HEKA) and current traces were visualized using IgorPro (v. 6.2.2.2, Wavemetrics, Lake Oswego, OR, USA). Collected data and statistical analysis were performed using GraphPad Prism (v. 8.4.3).

### Homology model for the extracellular domain binding site

The homology model of the ECD α_4_^(+)^β_1_^(-)^ interface has been described previously (40). The model was used to identify residues for the mutational study based on the docking of DS2 into the model described previously (40).

### Homology model for the transmembrane binding site

The homology model of the transmembrane part of the α_4_β_1_ interface was constructed with Modeller 9.24 (42) using the α_1_β_3_ interface from the α_1_β_3_γ_2L_ crystal structure (PDB code 6HUP (34)) as template. Model and template sequences of the TM helices and the connecting loops making up the subunit interface were obtained and aligned in UniProt (http://www.uniprot.org/ (43)); sequence IDs α_4_ P48169, β_1_ P18505, α_1_ P114867 and β_3_ P28472. To adhere as much as possible to the very closely related template structure, the “very fast” keyword was utilized to output the initial model that is only subjected to a brief optimization, thus, retaining the copied coordinates for all conserved residue positions. This procedure was selected based on the high sequence similarities and assumed structural conservation combined with the fact that the binding site residues are optimized relative to the ligand in following computational steps.

### Induced-fit docking of DS2 into the transmembrane α_4_^+^ β_1_^-^ site and in silico mutagenesis

The homology models were prepared for docking with the Protein Preparation Wizard (Schrödinger Release 2020-2, Schrödinger, LLC, New York, NY, 2020 (44)) using default settings. The chemical structure of DS2 was downloaded from the PubChem database (https://pubchem.ncbi.nlm.nih.gov (45) CID: 979718)) and the analogs DS2OMe (32) and DS2OPh (40) were built from DS2 in MarvinSketch 20.15.0, ChemAxon (http://www.chemaxon.com). All three ligands were prepared for docking with default settings in LigPrep (Schrödinger Release 2020-2) and used for induced-fit docking in the model of the transmembrane α_4_β_1_ interface with the Standard Protocol. The binding site center was defined by Ser303 and Ala324 from α_4_ plus Pro253 and Ile289 from β_1_. The ligand length was set to ≦ 14 Å and XP precision was used in the re-docking step, while all other settings were default. The best-scoring docking poses according to the IFD score were selected for each compound as the most likely binding mode. *In silico* mutagenesis was performed with the built-in protein mutagenesis wizard in PyMOL (Molecular Graphics System, Version 2.0 Schrödinger, LLC.) and the backbone dependent rotamer library selecting the most probable rotamer with the fewest steric clashes with surrounding residues. For α_4_S303L and β_1_I289Q mutations the first (1 of 4, 47.4%) and second (2 of 16, 14.6%) most likely rotamers were selected, respectively.

## Results and Discussion

To identify central residues for the activity of DS2 at δ-containing GABA_A_Rs, we systematically investigated three potential binding pockets: one in the ECD (the α_4_^(+)^δ^(-)^ interface), and two in the TMD (the α_4_^(+)^β_1_^(-)^ and the β_1_^(+)^α_4_^(-)^-interfaces). Although δ-containing TMD interfaces are present in the receptor complex we focused solely on the α-β interfaces due to the confirmed existence of binding sites at these interfaces (46–48), and our focus on benzodiazepine binding sites as potential binding sites for DS2 due to the structural resemblance between DS2 and zolpidem (40). Key interacting residues were identified using homology models and pharmacologically characterized in well-established HEK cell-based assays using the α_4_β_1_δ receptor as a model receptor, that has been carefully characterized in our hands (27, 28, 41).

## Investigation of the ECD α^(+)^-δ^(-)^ interface as a potential site of action for DS2

First, using the homology model published in (40), we studied the pocket located in the C-loop of the α^(+)^δ^(-)^ interface in the ECD of α_4_β_1_δ receptors (Fig. 1A). From our previous docking into the model, we identified three potential key residues on the α_4_^(+)^-side of the interface that could either interact directly with DS2 or were placed centrally within the binding pocket: α_4_(F133), α_4_(R135) and α_4_(G191) (Fig. 1A). Additionally, five residues on the complementary δ^(-)^-interface were identified: δ(E71), δ(A73), δ(F90), δ(S155) and δ(H204). The selected residues were mutated with the principle of removing potential interactions (into alanine) and/or gradually decreasing the space in the binding pocket (into various amino acid residues), thus expecting a reduced modulation by DS2 compared to WT. This resulted in seven different α_4_-subunit mutants and five different δ-subunit mutants (Table S1+S2). Each of the mutated subunits were expressed in HEK cell lines to form α_4_β_1_δ receptors and tested in the FMP assay, as single mutants. Whereas α_4_-mutants were simply co-transfected with WT β_1_ into WT stable δ-HEK293 Flp-In cells, each of the δ-mutants were established as stable HEK293 Flp-In cell lines transfected with WT α_4_β_1_ to transiently express α_4_β_1_δ. In general, extending the utility of this expression system from WT to mutated δ-containing receptors is highly reliable and suitable for controlling expression and reliably studying these in some instances, cumbersome receptor subtypes (49).

**Figure 1.**
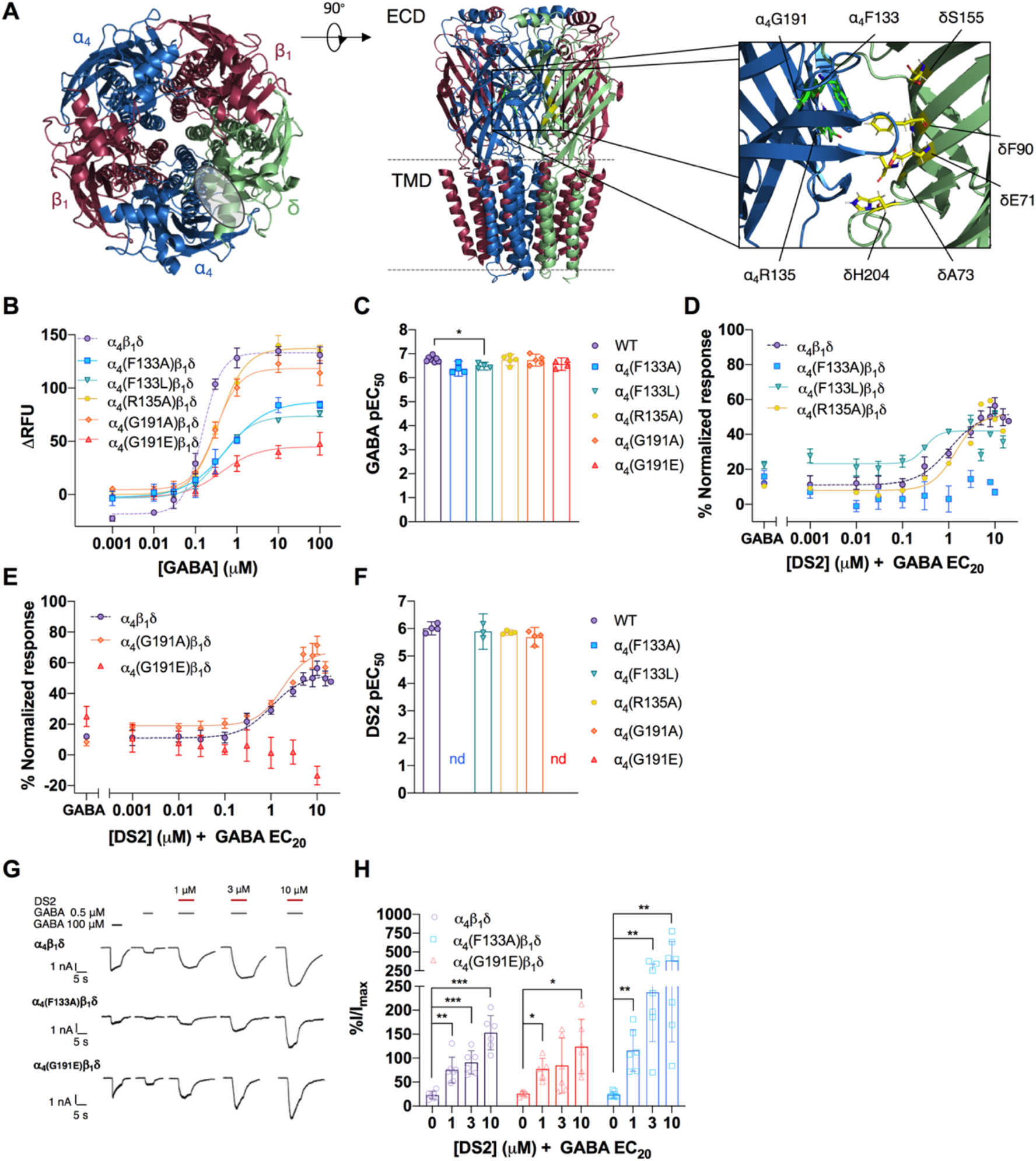
Investigation of the potential ECD α_4_^(+)^β^(-)^-interface binding site. **A** Model of the α_4_β_1_δ receptor with zoom-in on mutated residues in the α_4_^(+)^β^(-)^-interface (built on the cryo-EM structure of α_1_β_2_γ_2_ (PBD code: 6D6T). **B** Single representative GABA concentration-response curves for α_4_-mutant receptors (means±SD, technical triplicates), and **C** bar diagram of pooled pEC_50_ values (means with 95% CI, each point representing an independent replicate (n=4-8). **D-E** Concentration-response curves of the modulation of GABA EC_20_ by DS2 at α_4_-mutant receptors (normalized means±SD, technical triplicates), and **F** bar diagram of pooled pEC_50_ values (means with 95% CI, each point representing an independent replicate (n = 3-5)). **G** Single cell representative current traces from whole-cell patch-clamp electrophysiology recordings of the modulation of GABA EC_20_ induced currents by DS2 at WT α_4_β_1_δ and mutants α_4_(F133A)β_1_δ and α_4_(G191E)β_1_δ receptors. **H** Bar diagram summarizing the modulation by DS2 of α_4_ mutants α_4_(F133A), α_4_(G191E) cf. WT (note the broken y-axis). Currents were normalized to the maximum GABA current and presented as mean % I/I_max_ with 95% CI from minimum two independent transfections (n = 5-7). *For C,F,H:* Statistical analysis was performed using two-sided Welch’s t-test compared to WT (*C,F*) or control current (*H*) and adjusted for multiple testing using the Original FDR method of Benjamini and Hochberg with a discovery rate of 0.05. Statistical significance *P<0.05, **P<0.01, ***P<0.001.

All seven α_4_-mutant receptors were found to express functionally active receptors, and to respond to GABA with 2-3 times the potency observed for WT (Fig. 1B,C, Fig. S1, Table S1).

The expression levels of α_4_(F133A/L)β_1_δ and α_4_(G191E)β_1_δ appeared lower as compared to WT, as the max ΔRFU values were consistently reduced in all experiments (Fig. 1B). To characterize the sensitivity to DS2, it was applied together with a GABA EC_20_ concentration, calculated for each mutant (Table 1, S2). Among the seven different α_4_-mutants, α_4_(F133A)β_1_δ and α_4_(G191E)β_1_δ showed no apparent or only small modulation by DS2, whereas the potency of DS2 at the other α_4_-subunit mutants was either unchanged (α_4_(F133L)β_1_δ, α_4_(R135A) and α_4_(G191A/L)β_1_δ) or slightly increased, (α_4_(R135H)β_1_δ), compared to WT (Fig. 1D-F, Table 1+ Fig. S1, Table S2). Interestingly, only a single of the introduced mutations at α_4_(F133) and α_4_(G191) showed changed responses, which could not readily be explained.

**Table 1.** Potencies of DS2 at WT and ECD α_4_-mutant receptors determined in the FMP assay.

| Receptor | DS2 (PAM) | | 95% CI | difference pEC<sub>50</sub><br>[95% CI] | P-value | Adjusted p-value | EC<sub>20</sub><br>(μM) |
| --- | --- | --- | --- | --- | --- | --- | --- |
| | EC<sub>50</sub> (μM) | pEC<sub>50</sub>±SEM, n | | | | | |
| WT | 0.97 | (6.01 ± 0.076, 4) | 5.77;6.25 | - | - | - | 0.06 |
| $\alpha_4$(F133A)$\beta_1\delta$ | -<sup>a</sup> | n=5 | - | - | - | - | 0.12 |
| $\alpha_4$(F133L)$\beta_1\delta$ | 1.29 | (5.89 ± 0.153, 3) | 5.24;6.54 | -0.12 [-0.65;0.42] | 0.53 | 0.7857 | 0.12 |
| $\alpha_4$(R135A)$\beta_1\delta$ | 1.41 | (5.85 ± 0.031, 4) | 5.76;5.95 | -0.15 [-0.38;0.075] | 0.14 | 0.4500 | 0.06 |
| $\alpha_4$(G191A)$\beta_1\delta$ | 2.04 | (5.69 ± 0.111, 4) | 5.77;6.25 | -0.32 [-0.66;0.02] | 0.061 | 0.3050 | 0.12 |
| $\alpha_4$(G191E)$\beta_1\delta$ | -<sup>a</sup> | n = 5 | - | - | - | - | 0.06 |

<sup>a</sup> not able to fit, but a small potentiation seen for concentrations higher than 1 μM. EC<sub>20</sub>; calculated GABA concentration coapplied with the PAM. Statistical analysis; two-tailed Welch's t-test compared to WT, adjusted for multiple comparison using the original FDR (Benjamini and Hochberg) method with discovery rate of 0.05.

As we and others have previously observed methodological limitations in the FMP assay (26, 28), we suspected that the apparent lack of modulation could be due to sensitivity limitations. Thus, to follow up, the two mutants α_4_(F133A) and α_4_(G191E) were tested using whole-cell patch clamp electrophysiology. At both mutated receptors, DS2 modulated the GABA EC_20_-induced currents in a concentration-dependent manner similar to, or even more than, WT (Fig. 1G,H). An explanation for the discrepancy between the two assays could be the low expression of these specific mutant receptors which precludes the modulation to be picked up in the less sensitive FMP assay. Each of the five δ-subunit mutations were also tested in the FMP assay. These were all functional and displayed unchanged responsiveness to DS2 compared to WT (Fig. S2, Table S3,4).

Altogether, we conclude that the C-loop pocket in the ECD α_4_^(+)^δ^(-)^-interface is not the site responsible for the PAM effect of DS2. This is consistent with substituent and bio-isostere effects from a recent SAR study, focused on the ECD α_4_^(+)^δ^(-)^ as a potential binding site (40).

## Investigation of the TMD α_4_^(+)^β_1_^(-)^ interface as the site of modulation by DS2

Instead, we looked into two pockets in the TMD αβ-interfaces (specifically involving TM2), as potential recognition sites for DS2, which proved to be an advantageous avenue.

The first pocket is located in the β^(+)^α^(-)^ interface in a site equivalent to the recently identified low-affinity binding site for diazepam (46). We hypothesized analogous binding circumstances for diazepam and DS2 in the TMD. Mutations in α_4_β_1_δ TMD pockets were suggested based on the cryo-EM structure of the human GABA_A_R α_1_β_3_γ_2L_ (PDB-code: 6HUP) in combination with a sequence alignment due to the high (>90%) local sequence identity of the subunits within the TMD region of interest.

The mutations, β_1_(S290F) on the β_1_^(+)^ side and α_4_(L302Y) on the α4^(-)^ side were initially probed due to an apparent central positioning of the residues in the pocket and orientation towards diazepam in the cryo-EM structure (Fig. 2A). As a similar pocket is present at the reverse subunit interface, we also included the corresponding mutations in the α_4_^(+)^β_1_^(-)^ interface, α_4_(S303L) on the α_4_^(+)^ side and β_1_(I289Q) on the β_1_^(-)^ side. However, as this pocket appears noticeably smaller than the β_1_^(+)^α_4_^(-)^ pocket, these were mutated into more flexible and less bulky residues. Additionally, to probe both proposed pockets simultaneously, we included the double mutant receptors α_4_(L302Y,S303L)β_1_δ and α_4_β_1_(I289Q,S290F)δ. All of the introduced mutations were expected to revert hydrophilicity/hydrophobicity and introduce steric hindrance, and thus would be anticipated to decrease or altogether abolish the effect of DS2.

**Figure 2.**
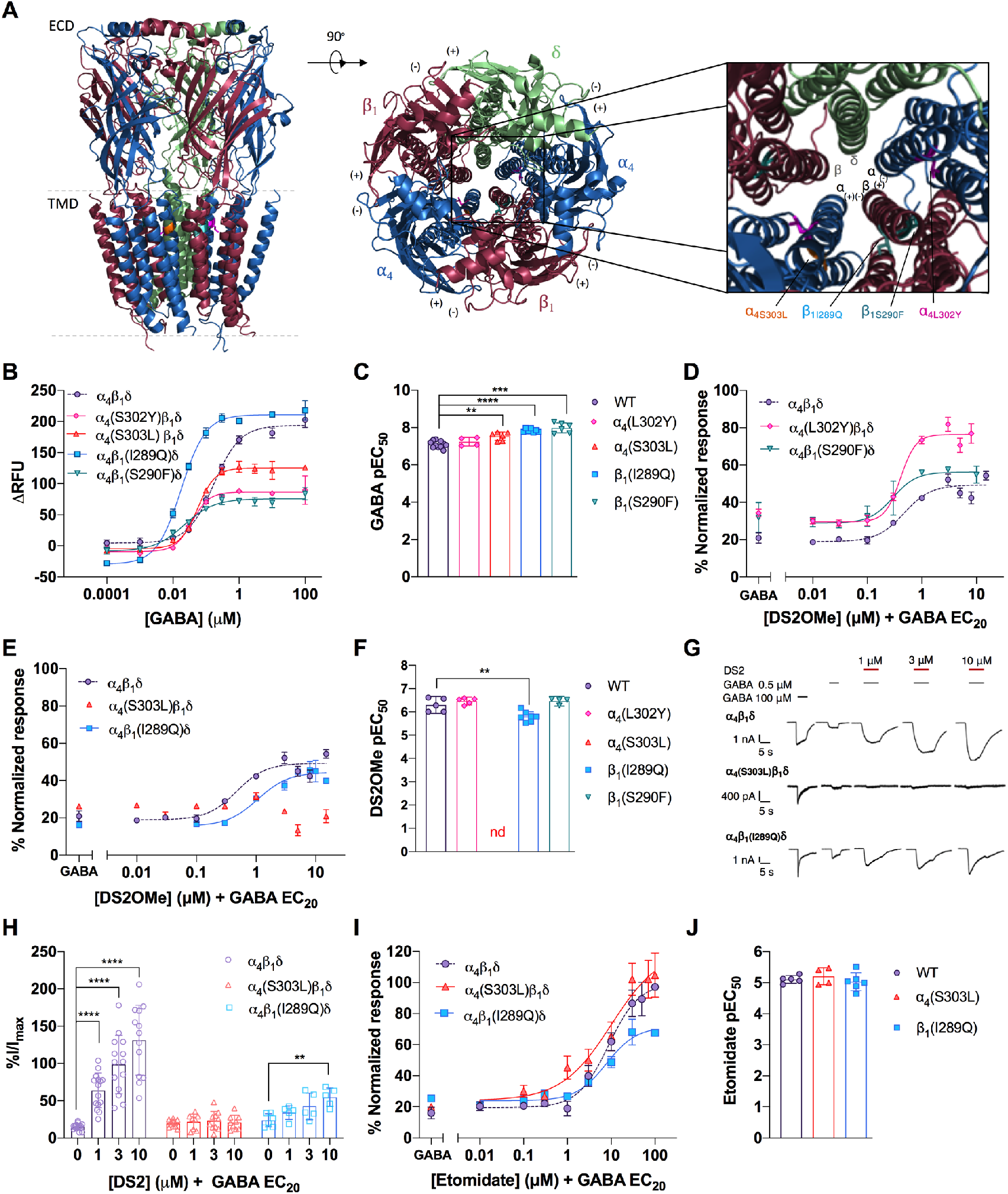
Identification of the TMD α_4_^(+)^β_1_^(-)^ site mediating modulation by DS2. **A** Model of the α_4_β_1_δ receptor with the TMD β_1_^(+)^α_4_^(-)^ and α_4_^(+)^β_1_^(-)^ interface mutants highlighted in the zoom-in (build on the cryo-EM structure of the α_1_β_3_γ_2_ receptor (PBD code: 6HUP). **B** Single representative GABA concentration-response curves from WT α_4_β_1_δ and TMD αβ interface mutants (means±SD, technical triplicates), and **C** bar diagram showing pooled pEC_50_ values (means with 95% CI (n=4-11). Modulation of GABA EC_20_ by DS2OMe at **D** the β_1_^(+)^α_4_^(-)^ and **E** the α_4_^(+)^β_1_^(-)^ interface mutants. Data are representative curves from a single experiment with means±SD of data normalized to GABA_max_. **F** Bar diagram showing pooled pEC_50_ values (means with 95% CI (n= 4-7). **G** Single cell current traces from DS2 modulation of GABA EC_20_ at WT and α_4_^(+)^β_1_^(-)^ interface mutants. **H** Bar diagram summarizing the DS2 modulation of α_4_^(+)^β_1_^(-)^ interface mutants in whole-cell patch-clamp electrophysiology. Currents are normalized to the GABA_max_ current and are given as mean % I/I_max_ with 95% CI (n= 5-11). **I** Single representative concentration-response curves of the modulation of GABA EC_20_ by etomidate at the α_4_^(+)^β_1_^(-)^ interface mutants, and **J** pooled pEC_50_ values (means with 95% CI with symbols representing values from independent experiments, (n = 4-5)). *For C,F,H,J:* Statistical analysis was performed using two-sided Welch’s t-test compared to WT (*C,F,J*) or control current (*H*) and adjusted for multiple testing using the Original FDR method of Benjamini and Hochberg with a discovery rate of 0.05. Statistical significance *P<0.05, **P<0.01, ***P<0.001 and ****P<0.0001.

First, we show that all the single mutant receptors were GABA-responsive and thus functional in the FMP assay (Fig. 2B). The two β-mutants β_1_(S290F) and β_1_(I289Q), displayed 6.8 and 8.1 times increased GABA potencies, respectively, and the α_4_(S303L)β_1_δ mutant 2.9 times increased potency compared to WT (Fig. 2C, Table S5). Since the receptors were functional, we continued with the studies.

In the modulation experiments, we switched to DS2OMe, an analog of DS2 (32) with the same pharmacological profile, because of both solubility issues with DS2 (described in the methods section) and the general sensitivity limitations observed in the FMP assay on the ECD mutants. First, we examined the modulation of GABA EC_20_ at mutations introduced in the β^(+)^α^(-)^ interface, equal to the low affinity diazepam binding site in the γ-containing receptor (Fig. 2D). These mutations did not affect the modulation by DS2OMe as both the α_4_β_1_(S29OF)δ and α_4_(L302Y)β_1_δ mutant receptors had DS2OMe potencies similar to WT, although a small significant increase in efficacy for the α_4_(L302Y)β_1_δ mutant compared to WT was observed (**P=0.0063, two-tailed Welch’s t-test, response of 3 μM DS2OMe) (Fig. 2D,F, Table 2). Altogether, we conclude that the β^(+)^α^(-)^ interface site is not involved in the effect of DS2, and that the changed potency of GABA did not affect the modulation of the receptors by DS2OMe.

**Table 2.**
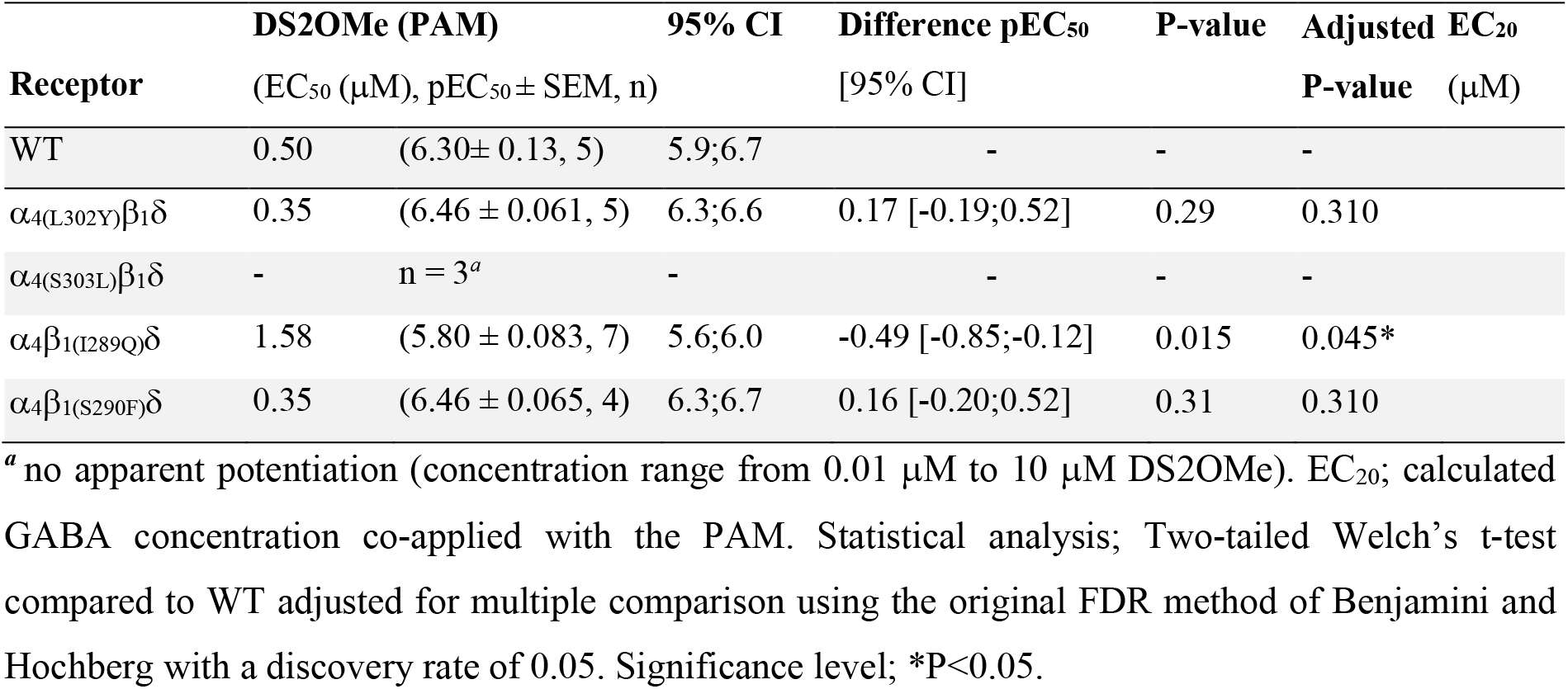
Potencies of DS2 at TMD mutant receptors determined in the FMP assay

By contrast, when turning to the alternative α_4_^(+)^β_1_^(-)^ interface, we observed significant decreases in responsiveness to modulation by DS2OMe. The α_4_(S303L)β_1_δ receptor lacked responsiveness to modulation by DS2OMe, and the β-mutant receptor, α_4_β_1_(I289Q)δ had a 3.2 times reduction of the potency of DS2OMe compared to the WT receptor (Fig. 2E,F, Table 2).

Additionally, as expected from the individual mutations, the double mutant receptor α_4_(L302Y,S303L)β_1_δ was not modulated by DS2OMe (Fig. S3, Table S6).

To verify the FMP results we performed whole-cell patch-clamp electrophysiology recordings. Convincingly, we found no or very limited DS2 modulation of the GABA currents in the α_4_(S303L)β_1_δ and α_4_β_1_(I289Q)δ receptor (only 10 μM modulated the α_4_β_1_(I289Q)δ receptor by significantly increasing the GABA control current to 54.8% of the GABA I_max_ (**P=0.0063, two-tailed Welch’s t-test, adjusted, n=5-6)) (Fig. 2G,H). Further, we included the double mutant receptor α_4_(S303L)β_1_(I289Q)δ, which was even less modulated by 10 μM DS2, amounting to 44% of the GABA I_max_ (*P=0.016, two-tailed Welch’s t-test, adjusted, n=6-9) (Fig. S4, Table S3,6). DS2OMe showed no modulation of the GABA response in either α_4_β_1_(I289Q)δ or α_4_(S303L)β_1_δ receptors, or the double mutant α_4_(S303L)β_1_(I289Q)δ receptor (Fig. S4,S5). Together, these results strongly advocate for the identified TMD α^(+)^β^(-)^ interface site as the site responsible for the modulatory action of DS2.

## Known GABA_A_R PAMs show unchanged modulation at DS2-insensitive mutant receptors

To confirm that the mutant receptors with altered DS2 sensitivity were not overall compromised in their general PAM responsiveness, we tested etomidate (50) and AA29504 (36, 51) at both the WT and the single mutants α_4_(S303L)β_1_δ and α_4_β_1_(I289Q)δ. In the FMP assay both compounds showed intact positive modulation of both mutants compared to WT. Potencies (EC_50_) of etomidate were at WT α_4_β_1_δ determined to 7.8 μM, and for the α_4_(S303L)β_1_δ and α_4_β_1_(I289Q)δ mutants to 6.2 μM and 9.3 μM, respectively (Fig. 2I,J, Table 3) (NS, two-tailed Welch’s t-test, n=3-4). Additionally, AA29504 showed similar potentiation at the mutants and WT receptors (Fig. S6, Table S7), indicating that it does not mediate its effect through the same site as DS2, correlating with a proposed binding site for AA29504 in the TMD β^(+)^α^(-)^ interface (36).

**Table 3.** Potency of etomidate at α_4_^(+)^β_1_^(-)^ TMD mutants determined in the FMP assay

| **Receptor** | **Etomidate (PAM)**<br>(EC<sub>50</sub> (μM), pEC<sub>50</sub> ± SEM, n) |  | **95% CI** | **Difference pEC<sub>50</sub>**<br>[95% CI] | **P-value** | **Adjusted P-value** | **EC<sub>20</sub>**<br>(μM) |
| --- | --- | --- | --- | --- | --- | --- | --- |
| WT | 7.8 | (5.11 ± 0.055, 5) | 5.1;7.4 | - | - | - | 0.06 |
| $\alpha_4(S303L)\beta_1\delta$ | 6.2 | (5.21 ± 0.14, 4) | 6.0;7.2 | -0.10 [-1.3;0.60] | 0.36 | 0.66 | 0.02 |
| $\alpha_4\beta_1(I289Q)\delta$ | 9.3 | (5.03 ± 0.12, 5) | 6.1;6.2 | -0.07 [-1.42;1.31] | 0.66 | 0.66 | 0.007 |

EC<sub>20</sub>; calculated GABA concentration coapplied with the PAM. Statistical analysis; two-tailed Welch's t-test compared to WT and adjusted for multiple testing using the original FDR method of Benjamini and Hochberg with a discovery rate of 0.05.

## Induced-fit docking of DS2 and DS2OMe corroborates mutational results

Guided by the mutational data confirming the molecular recognition site mediating the effect of DS2 and DS2OMe in the transmembrane part of the α_4_^(+)^β_1_^(-)^ subunit interface, we constructed a model of the modulator-receptor binding mode. Based on the structure of the desensitized α_1_β_3_γ_2L_ receptor bound to GABA and diazepam (34), we constructed a homology model of the α_4_β_1_ subunit interface into which DS2 and DS2OMe were fitted using induced-fit docking. Allowing residue side chains in the “empty” homology model to adapt to the modulators, we obtained very similar binding modes for the docked compounds (Fig. 3+S6). The core scaffold binds with the amide carbonyl of DS2 showing a potential hydrogen bond to the hydroxy group of Ser303 in α_4_. Using a backbone dependent rotamer library, we observe that the S303L mutation removes the hydrogen bond and sterically blocks the binding site – providing an explanation for the observed lack of potentiation on this mutant (Fig. 3B). Ile289 in β_1_ lines 4-fluorophenyl of DS2, contributing to the binding through substantial van der Waal contacts, and the I289Q mutation has a steric clash with DS2 (Fig. 3B). As for the α_4_(S303L) mutant, these effects provide a possible explanation for the observed abolishment of potentiation at all but the highest concentration of DS2 and DS2OMe in our patch clamp experiments and the obtained binding mode thus concurs with the experimental results. Our previously published analogs of DS2 show that there should be room for much larger substituents than the methoxy of DS2OMe as well as a bromo atom in the 5-position on the imidazol[1,2-a]pyridine scaffold (40). Thus, we provide further proof-of-concept for the predicted binding site by docking the recently published analog, DS2OPh (40). This confirmed that the OPh substituent can fit the binding site in the homology model with only a minor shift in the binding mode (Fig. S6).

**Figure 3.**
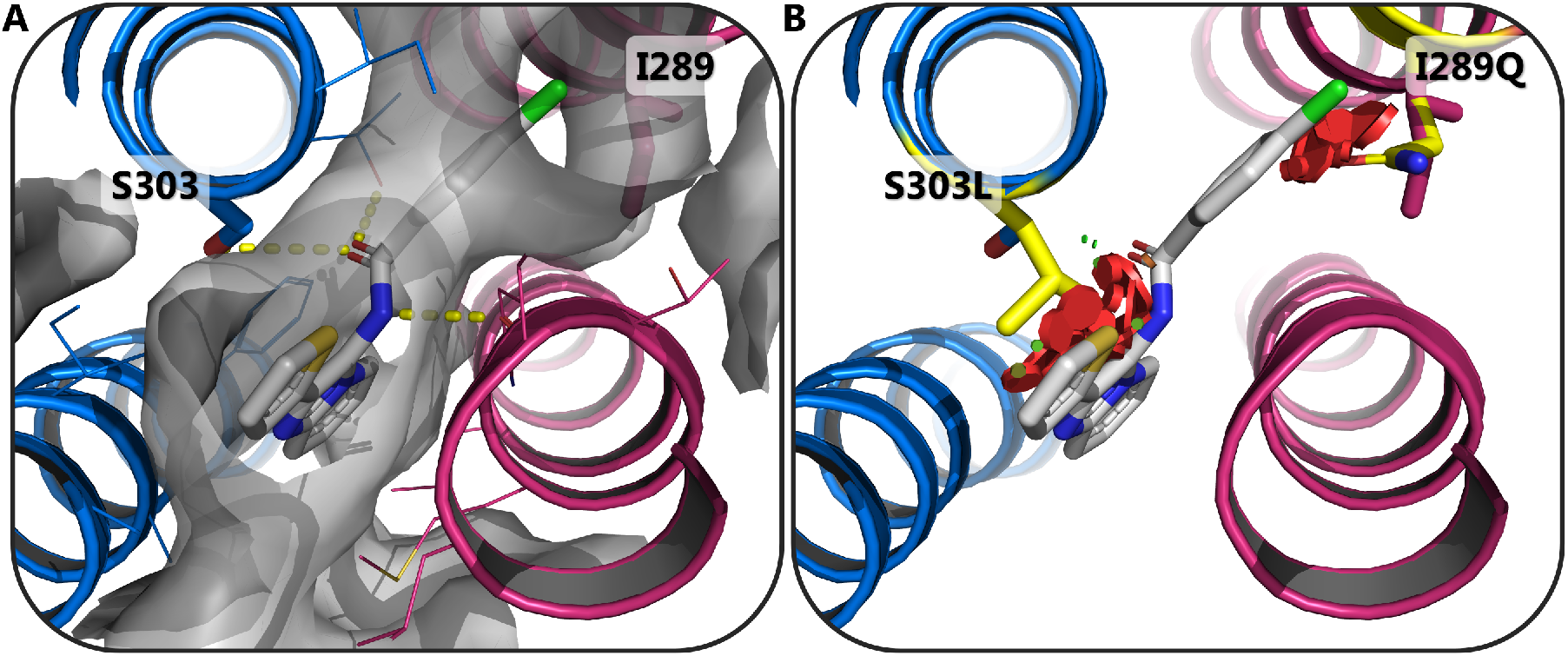
Binding model of DS2 in the TMD α_4_^(+)^β_1_^(-)^ subunit interface. Visualization of DS2 (sticks and grey carbon atoms) in the TMD interface between the α_4_ (blue cartoon and carbon atoms) and b_1_ (red cartoon and carbon atoms) GABA_A_ subunits, showing: **A** Residues with side chain atoms within 5 Å of DS2 are shown as lines highlighting the two important residues, α_4_S303 and β_1_I289, as sticks. The binding cavity is depicted as the vdW surface (grey and transparent) of the same residues and hydrogen bonds between DS2 and the receptor represented as yellow dotted lines. **B** *In silico* representation of the α_4_S303L and β_1_I289Q mutations (inserted residues as sticks with yellow carbon atoms) showing predicted steric clashes with DS2 as red disks explaining the hampered/abolished positive modulation of DS with these mutations. Figure prepared with the PyMOL Molecular Graphics System, Version 2.0 Schrödinger, LLC.

From our experiments using systematic structural iterations and experimental validation of known and proposed binding sites, we present the elusive DS2 interaction site encompassing α_4_S303 and β_1_I289 residues in the α_4_^(+)^β_1_^(-)^ interface. The proposed binding site for DS2 is distinct and novel, yet similar in nature to the low-affinity diazepam binding site (46) and the general anesthetics binding site located in the alternative interface. Indeed, it has been reported that pockets exist in all the TMD inter-subunit interfaces (37, 38), and that several known allosteric modulators can bind in these pockets (9). From our data it is deduced that the δ-subunit is not directly involved in the modulation by DS2, questioning what determines the δ-selective profile of the compounds. It is plausible that this is a matter of functional selectivity, similar to that observed for the super agonist THIP, in which case binding in a highly conserved α-β interface gives rise to 10-times higher potency at the δ-containing receptors compared to both γ-containing and binary αβ receptors with (16, 28, 52). Finally, having a homology model of the DS2 binding site, the next step is to use this for structure-based drug design of DS2-related analogs or a radiolabeled analog for further validation of the binding site.

Novel δ-selective analogs will aid to improve our understanding of the physiological and pathophysiological role of δ-containing receptors, and may, potentially, serve as leads for future rational drug development to treat the vast majority of neurological disorders with dysregulated tonic inhibition and/or target conditions involving δ-containing GABA_A_ receptors in the periphery.

## Author Contributions

C.B.F-P., F.R., K.H., D.E.G., B.F., and P.W. designed the research. C.B.F-P., F.R., R.L., S.B., and K.H. conducted the experiments. F.R. and B.F. provided new reagents. C.B.F-P., F.R., R.L., S.B., K.H. and P.W. analyzed data, and C.B.F-P., K.H., and P.W. wrote the paper.

## Competing interest statement

The authors declare no competing interests.

## Acknowledgements

We would like to thank Dr. Uffe Kristiansen for intellectual input and scientific guidance with the patch-clamp electrophysiology studies, and Durita Poulsen for help with performing DNA preps. This work was financially supported by the Lundbeck Foundation (grant R230-2016-2562 to C.B.F-P. and R277-2018-260 to P.W.), and the Drug Research Academy (C.B.F-P.). F.R was financially supported by a 2018 Lundbeck Foundation pre-graduate scholar stipend in Pharmaceutical Neuroscience.

## Notes

### Competing Interest Statement

The authors have declared no competing interest.

