## Supplementary Information for "Molecular determinants underlying DS2 activity at δ-containing GABA_A_ receptors"

**This PDF file includes:**

Figures S1 to S6
Tables S1 to S7

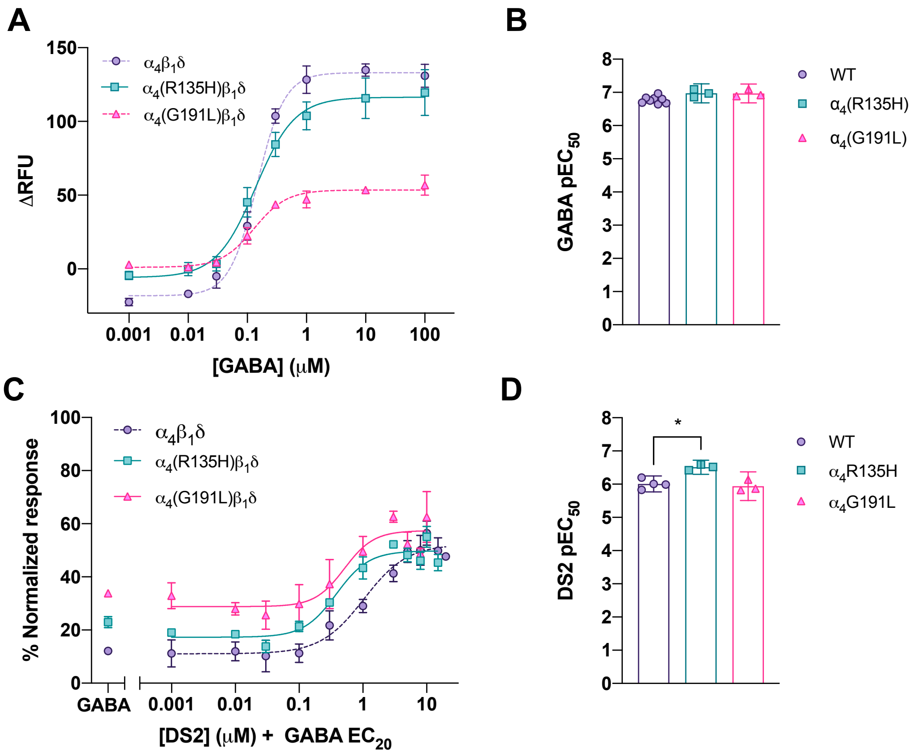


**Fig. S1.** Characterization of selected $\alpha_4$ ECD mutants in the FMP assay. **A** Representative GABA concentration-response curves shown as means±SD, and **B** bar diagram showing pooled GABA $pEC_{50}$ values (means with 95% CI, n=3-7). **C** Representative curves of the modulation of GABA $EC_{20}$ by DS2 shown as normalized means±SD and **D** bar diagram showing pooled $pEC_{50}$ values (means with 95% CI, n=3-4). Statistical analysis of $pEC_{50}$ values was performed using two-tailed Welch's t-test compared to WT and adjusted for multiple testing using the original FDR method of Benjamini and Hochberg with a discovery rate of 0.05. Statistical significance *$P<0.05$.

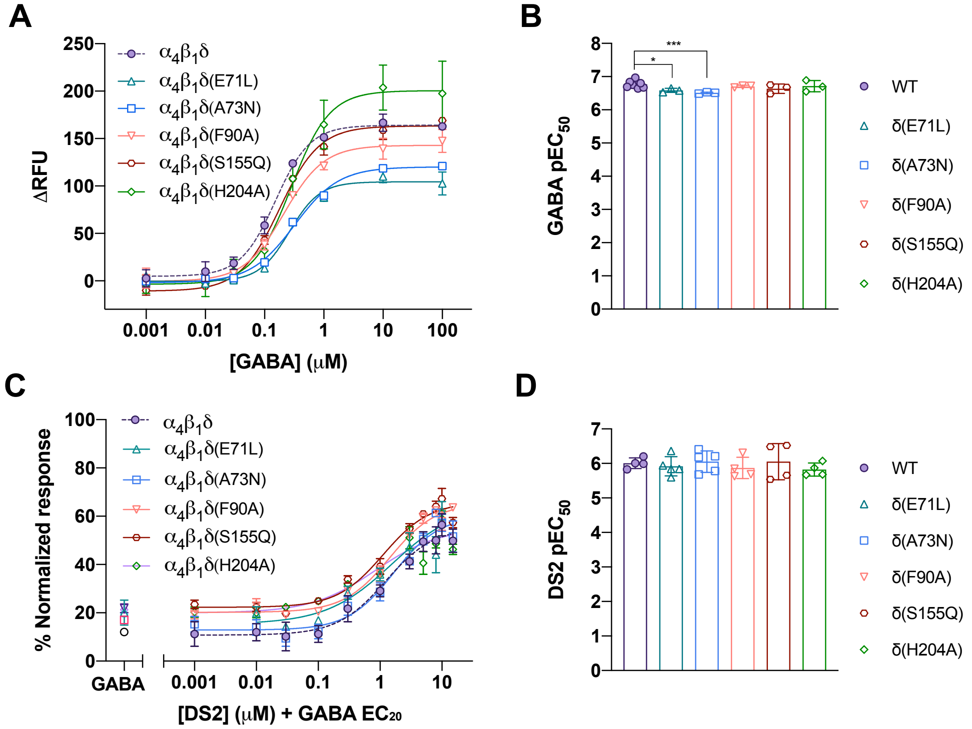


**Fig. S2.** Characterization of ECD δ-subunit mutant receptors in the FMP assay. **A** Representative GABA concentration-response curves shown as means±SD and **B** bar diagram showing pooled GABA $pEC_{50}$ values (means with 95% CI, n=3-8). **C** Representative curves of the modulation of GABA $EC_{20}$ by DS2 shown as normalized means±SD and **D** bar diagram showing pooled $pEC_{50}$ values (means with 95% CI, n=4-5). Statistical analysis of $pEC_{50}$ values was performed using two-tailed Welch's t-test compared to WT and adjusted for multiple testing using the original FDR method of Benjamini and Hochberg with a discovery rate of 0.05. Statistical significance *$P<0.05$ and ***$P<0.001$.

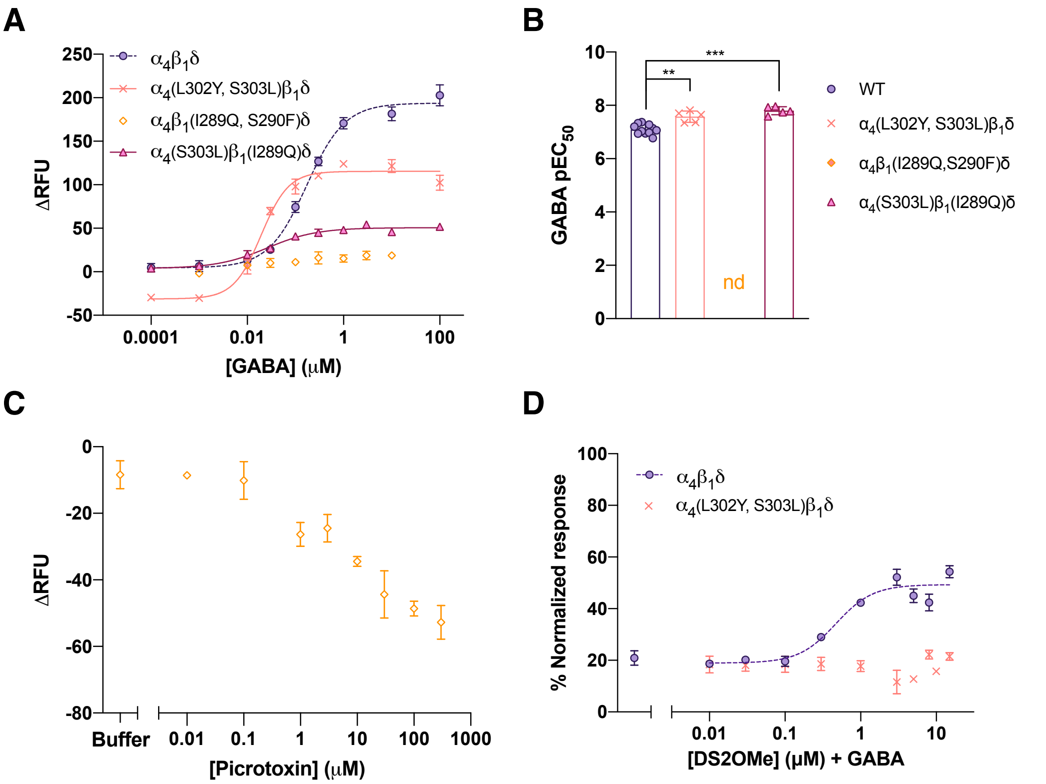


**Fig. S3.** Characterization of TMD double mutants in the FMP assay. **A** Representative GABA concentration-response curves (means±SD) at selected TMD double mutants and **B** bar diagram summarizing the pooled GABA $pEC_{50}$ values (means with 95% CI, n=5-11). The mutant $\alpha_4\beta_1$(I289Q,S290F)δ could not be fitted to a sigmoidal function due to low responses to GABA (n=3). **C** Decrease of the baseline signal by Picrotoxin at the $\alpha_4\beta_1$(I289Q,F290F)δ mutant, indicating constitutive activity (representative curve shown as normalized means±SD, n=2). **D** Representative curve of DS2OMe modulation at $\alpha_4$(L302Y,S303L)$\beta_1\delta$, means±SD (n=3). Statistical analysis of $pEC_{50}$ values was performed using two-tailed Welch's t-test compared to WT and adjusted for multiple testing using the original FDR method of Benjamini and Hochberg with a discovery rate of 0.05. Significance levels **P<0.01 and ***P<0.001.

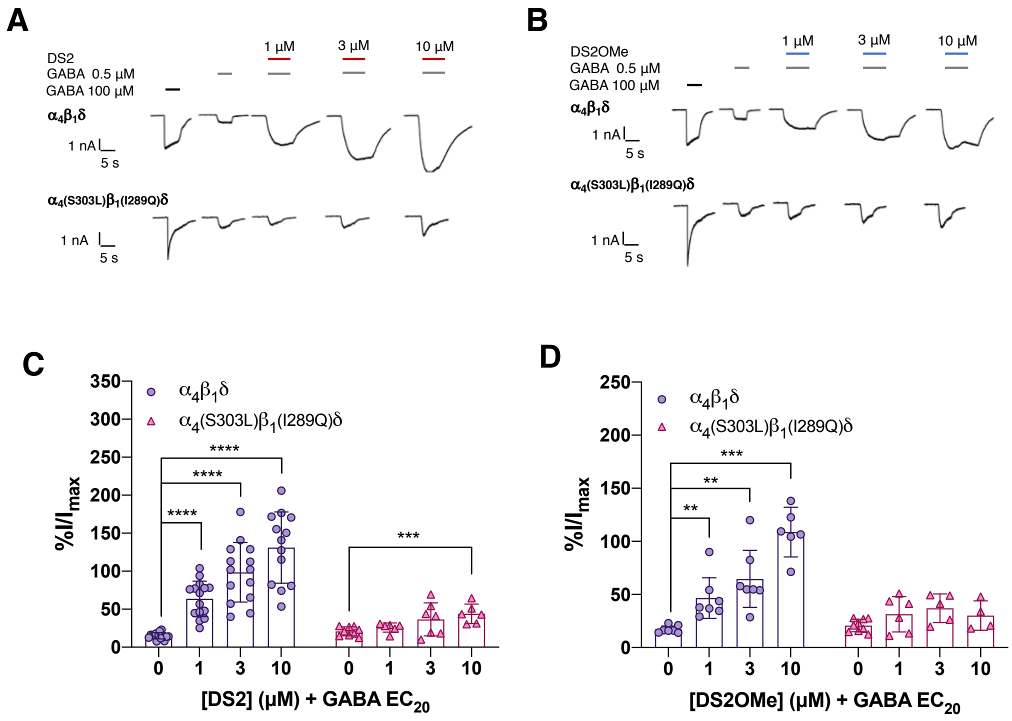


**Fig. S4.** Characterization of the TMD $\alpha_4^{(+)}\beta_1^{(-)}$ double mutant $\alpha_4$(S303L)$\beta_1$(I289Q)$\delta$ in whole cell patch clamp electrophysiology. **A** and **B** show representative cell current traces for the modulation of the double mutant receptor $\alpha_4$(S303L)$\beta_1$(I289Q)$\delta$ by DS2 and DS2OMe. In **C** and **D** are shown bar diagrams of the pooled normalized currents given as means with 95% CI from 5-16 cells. Statistical analysis was performed using two-tailed Welch's t-test compared to control current and adjusted for multiple testing using the original FDR method of Benjamini and Hochberg with a discovery rate of 0.05. Significance levels **P<0.01, ***P<0.001 and ****P<0.0001.

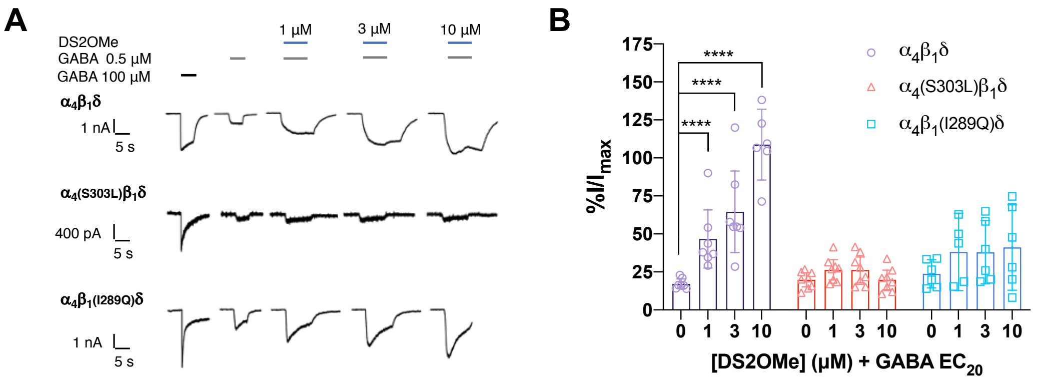


**Fig. S5**. **A** Representative cell current traces for DS2OMe modulation of WT $\alpha_4\beta_1\delta$ and $\alpha_4^{(+)}\beta_1^{(-)}$ interface mutant receptors. **B** Bar diagram summarizing the pooled currents normalized to the maximum GABA current shown as mean currents with 95% CI from 5-7 cells. Statistical analysis was performed using two-tailed Welch's t-test compared to control current and adjusted for multiple testing using the original FDR method of Benjamini and Hochberg with a discovery rate of 0.05. Significance level ****P<0.0001.

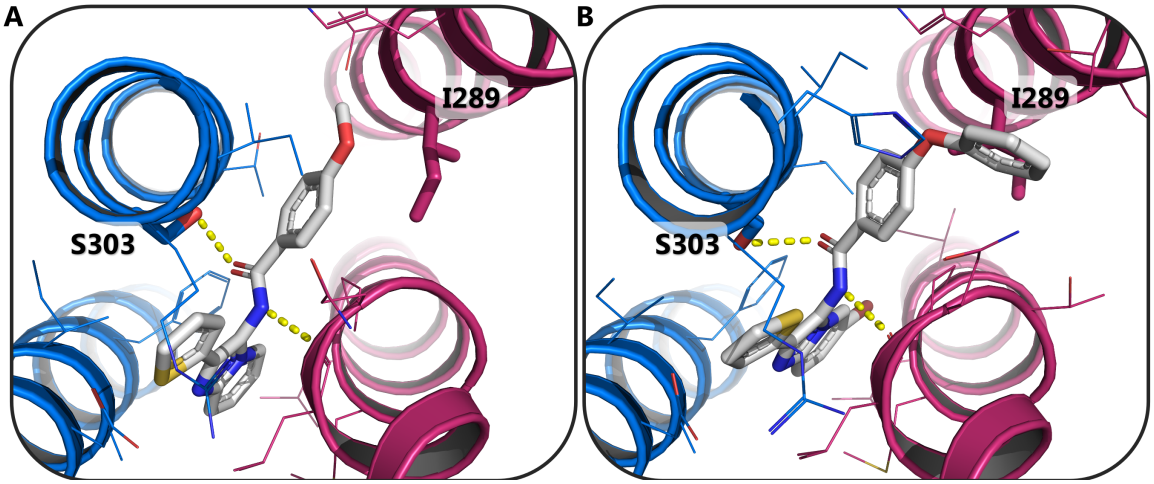


**Fig. S6.** Binding model of DS2OMe and DS2OPh in the TMD $\alpha_4\beta_1$ subunit interface. Potential binding mode of, **A**, DS2OMe and, **B**, DS2OPh as sticks and grey carbon atoms in the TMD interface between the $\alpha_4$ (blue cartoon and carbon atoms) and $\beta_1$ (red cartoon and carbon atoms) $GABA_A$ subunits. Residues with side chain atoms within 5 Å of the compounds are shown as lines highlighting the two important residues, $\alpha_4$S303 and $\beta_1$I289, as sticks. Hydrogen bonds to receptor are represented as yellow dotted lines. Figure prepared with the PyMOL Molecular Graphics System, Version 2.0 Schrödinger, LLC.

**Table S1.** GABA potencies at WT and ECD $\alpha_4$-mutant receptors.

| Receptor | **GABA (Agonist)** $EC_{50}$ (μM), $pEC_{50}$±SEM, n | | **95% CI** $pEC_{50}$ | **Difference $pEC_{50}$** mean [95% CI] | **P-value** | **Adjusted P-value** |
|---|---|---|---|---|---|---|
| WT | 0.17 | (6.77± 0.038, 8) | [6.68;6.86] | - | - | - |
| $\alpha_{4(F133A)}\beta_1\delta$ | 0.42 | (6.38 ± 0.097, 4) | [6.07;6.69] | -0.39 [-0.69;-0.10] | 0.020 | 0.071 |
| $\alpha_{4(F133L)}\beta_1\delta$ | 0.33 | (6.48 ± 0.091, 5) | [6.32;6.65] | -0.29 [-0.44;-0.14] | 0.0034 | 0.024* |
| $\alpha_{4(R135A)}\beta_1\delta$ | 0.18 | (6.74 ± 0.087, 5) | [6.50;7.98] | -0.03 [-0.27;0.20] | 0.75 | 0.75 |
| $\alpha_{4(R135H)}\beta_1\delta$ | 0.11 | (6.97 ± 0.066, 3) | [6.68;7.26] | -0.20 [-0.026;0.43] | 0.069 | 0.11 |
| $\alpha_{4(G191A)}\beta_1\delta$ | 0.18 | (6.74 ± 0.090, 5) | [6.48;6.98] | -0.04 [-0.28;0.21] | 0.71 | 0.75 |
| $\alpha_{4(G191E)}\beta_1\delta$ | 0.27 | (6.57 ± 0.082, 4) | [6.31;6.83] | -0.21 [-0.45;0.038] | 0.080 | 0.11 |
| $\alpha_{4(G191L)}\beta_1\delta$ | 0.18 | (6.97 ± 0.066, 4) | [6.69;7.25] | 0.20 [-0.026;0.42] | 0.069 | 0.11 |

Statistical analysis was performed using two-tailed Welch's t-test compared to WT and adjusted for multiple testing using the original FDR method of Benjamini and Hochberg with a discovery rate of 0.05. Significance level *P<0.05. Data was obtained in the FMP assay.

**Table S2.** Potencies of the PAM DS2 at WT and selected ECD $\alpha_4$-mutant receptors.

| Receptor | DS2 (PAM) EC50 (μM) pEC50±SEM, n | | 95% CI pEC50 | Difference pEC50 mean [95% CI] | P-value | Adjusted P-value | EC50 (μM) |
|---|---|---|---|---|---|---|---|
| WT | 0.97 | (6.01 ± 0.076, 4) | [5.8;6.3] | - | | - | 0.06 |
| $\alpha_{4(R135H)}\beta_1\delta$ | 0.31 | (6.51 ± 0.050, 3) | [6.3;6.7] | -0.50 [0.27;0.73] | 0.003 | 0.030* | 0.06 |
| $\alpha_{4(G191L)}\beta_1\delta$ | 1.15 | (5.94 ± 0.10, 3) | [5.5;6.4] | -0.07 [-0.41;0.28] | 0.63 | 0.79 | 0.06 |

$EC_{20}$; GABA concentration coapplied with the PAM. Statistical analysis; two-tailed Welch's t-test compared to WT, adjusted for multiple comparison using the original FDR (Benjamini and Hochberg) method with discovery rate of 0.05. $EC_{20;}$ modulated GABA concentration. Data was obtained in the FMP assay.

**Table S3.** GABA potencies at WT and ECD $\delta$-mutant receptors.

| Receptor | **GABA (Agonist)** EC$_{50}$ ($\mu$M), pEC$_{50}$±SEM, n | **95% CI** pEC$_{50}$ | **Difference pEC$_{50}$** mean [95% CI] | **P-value** | **Adjusted P-value** |
|---|---|---|---|---|---|
| WT | 0.17 (6.77± 0.038, 8) | [6.68;6.86] | - | - | - |
| $\alpha_4\beta_1\delta_{(E71L)}$ | 0.26 (6.59 ± 0.034, 3) | [6.44;6.74] | -0.18 [-0.31;-0.06] | 0.009 | 0.023* |
| $\alpha_4\beta_1\delta_{(A73N)}$ | 0.31 (6.51 ± 0.011, 3) | [6.46;6.56] | -0.26 [-0.35;-0.17] | 0.0002 | 0.0010*** |
| $\alpha_4\beta_1\delta_{(F90A)}$ | 0.19 (6.71 ± 0.018, 3) | [6.63;6.79] | -0.07 [-0.16;0.03] | 0.16 | 0.27 |
| $\alpha_4\beta_1\delta_{(S155Q)}$ | 0.23 (6.64 ± 0.079, 3) | [6.29;6.99] | -0.14 [-0.42;0.15] | 0.23 | 0.29 |
| $\alpha_4\beta_1\delta_{(H204A)}$ | 0.19 (6.71 ± 0.097, 3) | [6.29;7.14] | -0.06 [-0.42;0.30] | 0.62 | 0.62 |

Statistical analysis; two-tailed Welch's t-test compared to WT and adjusted for multiple testing using the original FDR method of Benjamini and Hochberg with a discovery rate of 0.05. Data was obtained in the FMP assay.

**Table S4**. DS2 potencies at WT and ECD $\delta$-mutant receptors.

| **Receptor** | **DS2 (PAM)** $EC_{50}$ ($\mu$M), $pEC_{50}$±SEM, n | **95% CI** $pEC_{50}$ | **Difference** $\mathbf{pEC_{50}}$ mean [95% CI] | **P-value** | **Adjusted P-value** | $\mathbf{EC_{20}}$ ($\mu$M) |
|---|---|---|---|---|---|---|
| WT | 0.97 (6.01 ± 0.076, 4) | [5.8;6.3] | - | - | - | 0.06 |
| $\alpha_4\beta_1\delta_{(E71L)}$ | 1.20 (5.92 ± 0.13, 5) | [5.6;6.3] | -0.09 [-0.44;0.26] | 0.55 | 0.79 | 0.06 |
| $\alpha_4\beta_1\delta_{(A73N)}$ | 1.10 (5.96 ± 0.14, 4) | [5.7;6.4] | 0.04 [-0.34;0.43] | 0.79 | 0.87 | 0.06 |
| $\alpha_4\beta_1\delta_{(F90A)}$ | 1.35 (5.87 ± 0.15, 4) | [5.4;6.4] | -0.13 [-0.60;0.32] | 0.46 | 0.79 | 0.06 |
| $\alpha_4\beta_1\delta_{(S155Q)}$ | 0.89 (6.05 ± 0.26, 4) | [5.2;6.9] | -0.05 [-0.76;0.85] | 0.87 | 0.87 | 0.06 |
| $\alpha_4\beta_1\delta_{(H204A)}$ | 1.51 (5.82 ± 0.096, 4) | [5.5;6.1] | -0.19 [-0.48;0.11] | 0.18 | 0.45 | 0.06 |

$EC_{20}$, GABA concentration coapplied with the PAM. Statistical analysis; two-tailed Welch's t-test, adjusted for multiple comparison using the original FDR method of Benjamini and Hochberg with a discovery rate of 0.05. Data is from the FMP assay.

**Table S5.** GABA potencies at WT and TMD mutant receptors.

| Receptor | **GABA (Agonist)** $EC_{50}$ (μM), $pEC_{50}$ ± SEM, n | | **95% CI** $pEC_{50}$ | **Difference $pEC_{50}$** mean [95% CI] | **P-value** | **Adjusted P-value** |
|---|---|---|---|---|---|---|
| WT | 0.081 | (7.09 ± 0.056, 11) | [7.0;7.2] | - | - | - |
| $\alpha_{4(L302Y)}\beta_1\delta$ | 0.058 | (7.24 ± 0.12, 4) | [6.9; 7.6] | 0.14 [-0.20;0.49] | 0.31 | 0.3100 |
| $\alpha_{4(S303L)}\beta_1\delta$ | 0.029 | (7.54 ± 0.091, 6) | [7.3; 7.8] | 0.44 [0.20;0.69] | 0.0024 | 0.0032** |
| $\alpha_4\beta_{1(I289Q)}\delta$ | 0.010 | (7.84 ± 0.037, 7) | [7.8; 7.9] | 0.74 [0.60;0.89] | <0.0001 | 0.0002*** |
| $\alpha_4\beta_{1(S290F)}\delta$ | 0.012 | (7.98 ± 0.11, 6) | [7.7; 8.3] | 0.87 [0.60;1.17] | 0.0001 | 0.0002*** |

Statistical analysis; two-tailed Welch's t-test, adjusted for multiple comparison using the original FDR method of Benjamini and Hochberg with a discovery rate of 0.05. $EC_{20}$ is the used GABA concentration. Data is from the FMP assay.

**Table S6**. GABA potencies at WT and TMD $\alpha 4^{(+)}\beta 1^{(-)}$ interface double mutant receptors.

| Receptor | GABA (agonist) EC50 (μM), pEC50 ± SEM, n | | 95% CI pEC50 | Difference pEC50 mean [95% CI] | P-value |
|---|---|---|---|---|---|
| WT | 0.081 | (7.09±0.056, 11) | [7.0;7.2] | - | - |
| $\alpha_{4(L302Y,S303L)}\beta_1\delta$ | 0.027 | (7.57 ± 0.094, 5) | [7.3;7.8] | 0.48 [0.22;0.74] | 0.003** |
| $\alpha_4\beta_{1(I289Q,S290F)}\delta$ | - | n = 3[a] | - | - | - |
| $\alpha_{4(S303L)}\beta_{1(I290Q)}\delta$ | 0.016 | (7.80 ± 0.067, 5) | [7.6;8.0] | 0.70 [0.50;0.90] | >0.0001*** |

[a] No response to GABA. Statistical analysis was performed using two-tailed Welch's t-test. Data was obtained in the FMP assay.

**Table S7.** AA29504 potencies at TMD α4(+)β1(-) interface mutants.

| Receptor | **AA29504 (PAM)** EC$_{50}$(μM), pEC$_{50}$ ± SEM, n | **95% CI** pEC$_{50}$ | **Difference pEC$_{50}$** mean [95% CI] | **P-value** | **EC$_{20}$** (μM) |
|---|---|---|---|---|---|
| WT | 0.55 (6.26 ± 0.27, 3) | [5.0;5.3] | - | - | 0.06 |
| $\alpha_{4(S303L)}\beta_1\delta$ | 0.25 (6.61 ± 0.20, 4) | [4.8;5.6] | -0.35 [-0.31;0.51] | 0.53 | 0.02 |
| $\alpha_4\beta_{1(I289Q)}\delta$ | 0.76 (6.12 ± 0.015, 3) | [4.7;5.3] | 0.14 [-0.38;0.23] | 0.58 | 0.007 |

EC$_{20}$; GABA concentration coapplied with the PAM. Statistical analysis was performed using two-tailed Welch's t-test. Data was obtained in the FMP assay.
